# The foraging brain: evidence of Lévy dynamics in brain networks

**DOI:** 10.1101/041202

**Authors:** Tommaso Costa, Giuseppe Boccignone, Franco Cauda, Mario Ferraro

## Abstract

In this research we have analyzed functional magnetic resonance imaging (fMRI) signals of different networks in the brain under resting state condition.

To such end, the dynamics of signal variation, have been conceived as a stochastic motion, namely it has been modelled through a generalized Langevin stochastic differential equation, which combines a deterministic drift component with a stochastic component where the Gaussian noise source has been replaced with *α*-stable noise.

The parameters of the deterministic and stochastic parts of the model have been fitted from fluctuating data. Results show that the deterministic part is characterized by a simple, linear decreasing trend, and, most important, the *α*-stable noise, at varying characteristic index *α*, is the source of a spectrum of activity modes across the networks, from those originated by classic Gaussian noise (*α* = 2), to longer tailed behaviors generated by the more general Lévy noise (1 ≤ *α* < 2).

Lévy motion is a specific instance of scale-free behavior, it is a source of anomalous diffusion and it has been related to many aspects of human cognition, such as information foraging through memory retrieval or visual exploration.

Finally, some conclusions have been drawn on the functional significance of the dynamics corresponding to different *α* values.

**Author Summary:** It has been argued, in the literature, that to gain intuition of brain fluctuations one can conceive brain activity as the motion of a random walker or, in the continuous limit, of a diffusing macroscopic particle.

In this work we have substantiated such metaphor by modelling the dynamics of the fMRI signal of different brain regions, gathered under resting state condition, via a Langevin-like stochastic equation of motion where we have replaced the white Gaussian noise source with the more general *α*-stable noise.

This way we have been able to show the existence of a spectrum of modes of activity in brain areas. Such modes can be related to the kind of “noise” driving the Langevin equation in a specific region. Further, such modes can be parsimoniously distinguished through the stable characteristic index *α*, from Gaussian noise (*α* = 2) to a range of sharply peaked, long tailed behaviors generated by Lévy noise (1 ≤ *α* < 2).

Interestingly enough, random walkers undergoing Lévy motion have been widely used to model the foraging behaviour of a range of animal species and, remarkably, Lévy motion patterns have been related to many aspects of human cognition.

## Introduction

Spontaneous brain activity is not random [1] and brain fluctuations have statistical properties that can be useful to characterize the state space within which brain dynamics evolves [2]. To shed light on such issue, in this work we have analyzed fMRI signals of different regions in the brain under resting state condition.

Comparison of BOLD signals amplitude in different conditions via univariate [3] or multivariate statistics [4-6] highlights the functional properties of a brain region, while other approaches aim to investigate such properties via methods of time series analysis such as linear correlation [7], Granger causality [8] and statistical modelling [9].

More relevant for the work reported here, a different line of research has demonstrated the importance of the variance of the BOLD signal to provide information about the working of the brain. For example it has been shown that signal variance decreases in the visual cortex during a task [10] and that aging correlates with decreased signal variance [11]. An in-depth study by [12] showed that both the variance and the power-law exponent of the BOLD signal decrease during task activation suggesting that the signal exhibits longer range memory during rest than during task. Thus, the overall brain signal variability across large-scale brain regions has emerged as a marker of a well-functioning brain [13]. Finally, in [14] it has been proved that age-related dopamine losses contribute to changes in brain signal variability.

Yet, when quantitative measurements of BOLD signal variability have been made, a deceptively simple question has been overlooked: what kind of dynamics is behind such variability and how is modulated in different networks?

Consider, for instance, Fig. 1 showing the detrended traces of the BOLD signal for the basal ganglia (red trace) and the cerebellum I (blue trace) respectively. (see Methods Section below).

**Figure 1.**
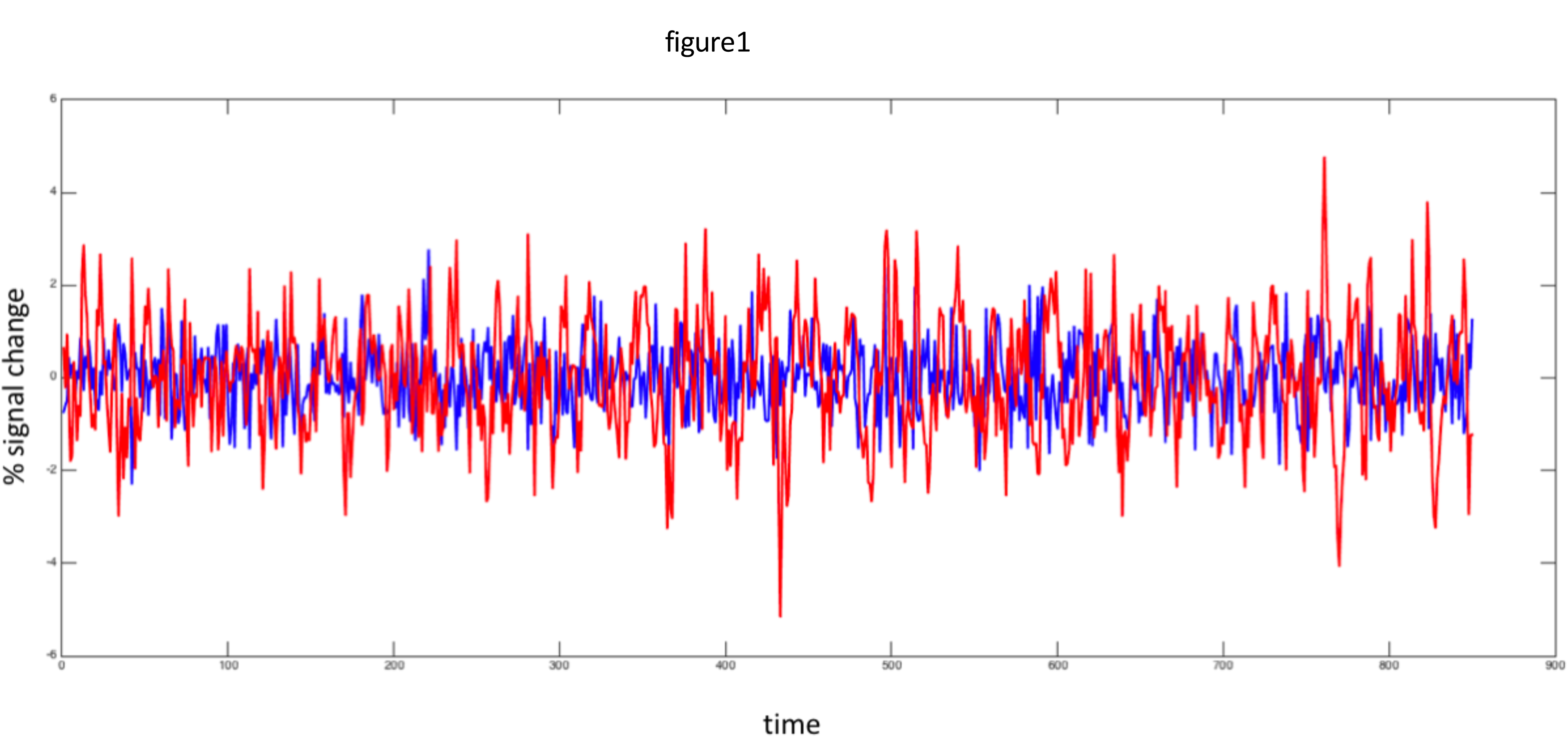
Examples of BOLD time series for two different networks. The traces are the detrended BOLD signals from the basal ganglia (red) and cerebellum I (blue) respectively.

It can be appreciated at a glance that signals exhibit a very different behavior. As opposed to the cerebellum I signal activity, which is confined within a roughly constant interval embedding small/average fluctuations, the signal from the basal ganglia exhibits a large number of tiny fluctuations interspersed with occasionally long “jumps”. Accounting for these differences is precisely the purpose of this paper.

We propose a unified model of the stochastic process underlying the different fMRI signal time courses from different brain regions. To such end we resort to a generalized Langevin stochastic differential equation. The Langevin equation is apt to decompose the signal evolution as the sum of a deterministic component (or drift) and a stochastic component responsible for noise (diffusion) [15,16]. Here, we replace the white Gaussian noise, which is normally used as the stochastic source, with the more general *α*-stable noise [17].

The main result of this study is that such model, when the forms of both components have been derived in a data-driven way, allows to precisely characterize a spectrum of activity of brain areas, the latter related to the kind of noise source governing the stochastic component specific to the area. Such spectrum ranges from the mode originated by adopting classic Gaussian white noise to longer tailed behaviors characterizing Lévy noise [17] generated from *α*-stable distributions. The latter are a specific type of the general family of stable Lévy processes that have very interesting properties for describing the complex behaviour of non-equilibrium dissipative systems such as turbulence and anomalous diffusion [18], [19]. Stable Lévy processes are infinitely divisible random processes, and they have the property of scaling (self-similarity), meaning that the variable of interest can be studied through a stable distribution of exactly the same form for each level of scale.

It is well known that the brain is the source of fluctuations with complex scaling properties (see [2] for an in-depth discussion) and indeed scale-free and power law properties are of clear interest for fMRI signal analysis, as mentioned before [12]. Remarkably, the interest for stable Lévy processes has moved beyond the frontiers of physics reaching a great number of distinct fields and studies concerning seismic series and earthquakes [20], spreading of epidemic processes [21], time series analysis of DNA [22], animal foraging such as albatross flights [23]. More relevant for the work presented here is that they have gained currency in the modelling of human memory retrieval [24] and eye movements behavior [25-28]. Thus, by adopting the theoretical model of a generalized Langevin equation accounting for Lévy motion, connections to different theoretic frameworks can be easily established [29]. For instance, it provides a viable approach to handle in a unique modelling framework either animal foraging and cognitive foraging, which substantiates Hills’ adaptation hypothesis [30] that what was once foraging for tangible resources in a physical space became, over evolutionary time, foraging in cognitive space for information related to those resources. Also, it gives a clear form to the intuition that brain fluctuations can be understood by conceiving brain activity as the motion of a random walker or, in the continuous limit, of a diffusing macroscopic particle [2].

At the best of our knowledge, no previous work has considered the important and wide class of dynamical Lévy systems for analysing fMRI time series in different brain areas.

## Methods

### Participants

Twenty-five healthy right-handed adults with no prior history of neurological or psychiatric impairment, participated in this study. All participants gave written informed consent for the study, approved by the University Ethics Board.

### Scanning

Subjects were instructed simply to keep their eyes closed, think of nothing in particular, and not to fall asleep. After the scanning session, participants were asked if they had fallen asleep during the scan, and data from subject with positive or doubtful answers were excluded from the study.

### Data acquisition

Images were gathered on a 1.5 T INTERA™scanner (PhilipsMedical Systems) with a SENSE high-field, high resolution (MRIDC) head coil optimized for functional imaging. Two resting state functional T2* weighted runs were acquired using echoplanar (EPI) sequences, with a repetition time (TR) of 2000ms, an echo time (TE) of 50 ms, and a 90° flip angle. The acquisition matrix was 64 × 64, with a 200 mm field of view (FoV). A total of 850 volumes were acquired for the first run and a total of 450 volumes were acquired for the second, each volume consisting of 19 axial slices, parallel to the anterior posterior (AC-PC) commissure; slice thickness was 4.5 mm with a 0.5 mm gap. To reach a steady-state magnetisation before acquiring the experimental data, two scans were added at the beginning of functional scanning: the data from these scans were discarded. Within a single session for each participant, a set of three dimensional high-resolution T1-weighted structural images was acquired, using a Fast Field Echo (FFE) sequence, with a 25 ms TR, an ultrashort TE, and a 30° flip angle. The acquisition matrix was 256 × 256, and the FoV was 256 mm. The set consisted of 160 contiguous sagittal images covering thewhole brain. In-plane resolution was 1 mm × 1 mm and slice thickness 1 mm (1 × 1 × 1 mm^3^ voxels).

### Data preprocessing

Analyses of the FMRI data were performed using the validated software package FSL (available from the FMRIB Software Library at www.fmrib.ox.ac.uk/fsl). Data were corrected for motion artefact, compensating for any head movements using an FSL linear (affine) transformation (FSL-MCFLIRT) procedure. Extraction of functional data from the brain scan was performed using the FSL brain extraction tool (FSL-BET). Functional data were spatially smoothed using a Gaussian kernel of 8 mm full width at half maximum. High-pass temporal filtering (50 s) was also performed, since we wanted to remove very low frequency scanner-drift artefacts.

Analyses of the resting-state FMRI data were performed using ICA with the FSL Multivariate Exploratory Linear Optimized Decomposition into Independent Components (FSL-MELODIC) tool and the validated dual-regression approach. These techniques allow for voxel-wise comparisons of resting-state functional connectivity by, first, temporally concatenating resting-state FMRI data from all subjects, followed by back-reconstructing the group ICNs for individual subjects, yielding data that can then be used for within-subject and between-subject group difference maps. This technique has been found to have moderate-to-high test-retest reliability in previous studies. Functional data were first projected into native anatomical space using Boundary-Based Registration (BBR) approach. Then native T1 scan was registered to standard Montreal Neurological Institute space using structural linear (affine) coregistration (FSL-FLIRT). These BOLD functional data were then concatenated in time across all subjects, creating a single 4-dimensional (4-D) data set. We then applied probabilistic ICA (with the FSL-MELODIC tool) to identify global, distinct (independent) patterns of functional connectivity in the entire subject population. We limited the number of independent components (ICs) in this step to 35. From this pool of 35 ICs, ICNs of interest were selected by visual inspection and we selected 11 ICA components as functional significant networks.

**Figure 2.**
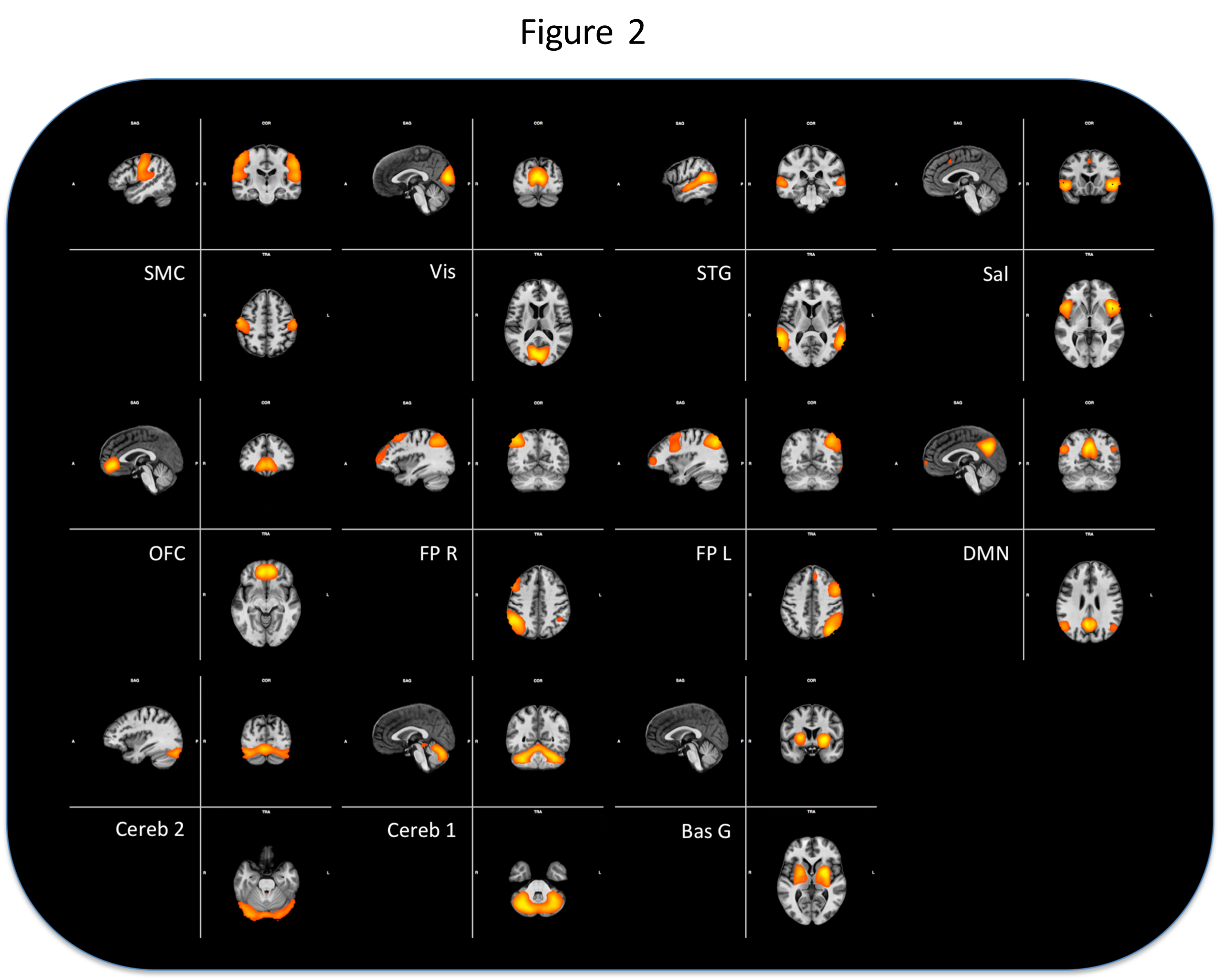
Networks used in the analisys. The 11 networks obtained with the ICA decomposition, see text for explanation.

In conclusion, we have obtained BOLD activity from a sample of 25 healthy human subjects, during sessions of 850 s, with *TR* = 2 s, in resting state condition. The dataset thus consists of 11 time series representing the networks obtained by a dual regression of the preprocessed resting state for each subject.

## Analysis

Consider a given network: in the following *x*(*t*) will denote its activity at time *t* as measured from the BOLD signal and the activity variation is written as

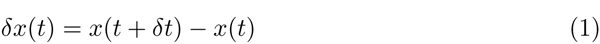

*δt* being the sampling time.

A useful framework to describe the process underlying the measured signal is provided by a random motion model, in which *x*(*t*) is the position in the one-dimensional activity space at time *t*. The use of the random motion metaphor in computational neuroscience indeed goes a long way back, see for instance [31, 32]; more recent examples can be found in [33, 34].

This framework suggests then a suitable mathematical representation of the process in terms of a stochastic differential equation, namely the Langevin equation in the coordinate *x*, which describes the evolution of *x* as driven by the sum of a deterministic and a stochastic component, commonly referred to as drift and diffusion, respectively [15,35, 36]

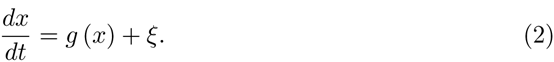

Here *g* represents the deterministic part of the process, whereas stochastic factors are described by the random variable (RV) *ξ*; as usually done in the literature, it will be assumed that 〈*ξ*〉 = 0, where 〈·〉 denotes the mean or expectation value of a RV. This implies that

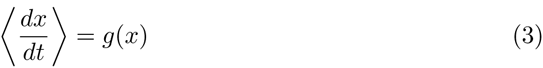

Now consider *ξ* here we refer again to the theory of random motions and note that different types of dynamics can be generated by resorting to the class of the so called *α*-stable distributions. Indeed the use of *α*-stable distributions is common to most applications of random walk and motions, the reason being that this is a very general class including distributions that can describe basically different types of processes (motions) from classical Brownian motion to Lévy flights [37], [18]

These distributions form a four-parameter family of continuous probability densities functions (pdf), say *ƒ*(*x;α,β,γ,δ*). The different parameters are, respectively, the skewness *β* ∈ [−1; 1] (a measure of asymmetry), the scale *γ* > 0 (width of the distribution), the location *δ* ∈ ℝ and, most importantly for our purposes, the index of the distribution (or characteristic exponent) *α*, that specifies the asymptotic behavior of the distribution [37], [18].

The relevance of *α* resides in the fact that [18,19] the probability density function p(|*ξ*|) of absolute values of *ξ* scales, asympotically, as

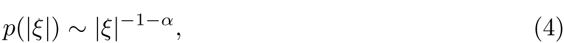

with *α* > 0 to ensure normalization, and this property defines two basically different types of random processes. If *α* ≥ 2 the pdf *ƒ* belongs to the domain of attraction of the Gaussian distribution, in the sense of the Central Limit Theorem: the sum of i.i.d random variables with pdf *ƒ*, follows, asymptotically, a Gaussian distribution. On the contrary if *α* < 2, then *ƒ*, the probability density function is characterized by heavy tails, meaning that the occurrence of large fluctuations is more likely than in Gaussian case [18]. In random walk theory the case with *α* ≥ 2 corresponds to the usual Brownian motion, whereas *α* < 2 gives rise to Lévy motion, a type of superdiffusive random motion, in which long jumps (displacements) are more likely. Thus it is clear that the value of the parameter *α* provides relevant information on the type of random process the activity of a given network can be associated with, allowing a discrimination between Gaussian and Lévy-like modes of activity.

The next step in our analysis is then to verify, for each network under consideration, that *ξ* indeed follow an *α*-stable distribution and to calculate the corresponding value of *α*. This is done, usually, by considering the upper tail of the survival function 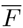(|*ξ*|), which is the complement of the cumulative distribution function *F*(|*ξ*|). A rationale is as follows.

Let *F* be the cumulative distribution of | *ξ* |, and let |*ξ*_p_| large enough so that for |*ξ*| ≥ |*ξ*_p_| the condition (4) holds.

Obviously,

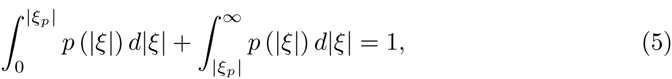

from which

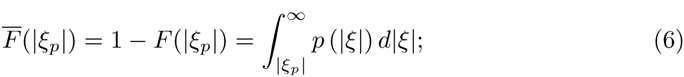

from condition (4) it follows that, for |*ξ*| ≥ |*ξ*_p_|,

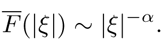

In conclusion then, if *F*(|*ξ*|) is *α*-stable, the graph of log 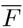 should exhibit a linear tail whose slope is just −*α*.

For future purposes, it is useful to note that the value of *α* does not depend on the sampling time *δt*. Let *k* > 0 be a constant, define *y* = *kξ* and let *τ* be the pdf of |*y*|. Suppose that condition (4) holds for *ξ*.

Probability conservation requires

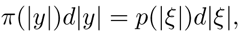

therefore

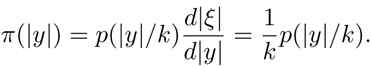

Asymptotically

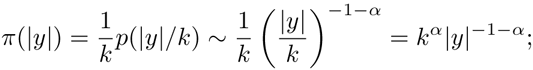

*k* is a constant, therefore

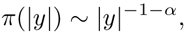

as it should be expected from the scale invariance of monomial functions. In other words pdfs of |*ξ*| and |*y*| share the same asymptotic trend, as far as *α* is concerned. Denote by 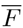(|*ξ*|) and Ḡ(|*y*|) the survival function of |*ξ*| |*y*|, respectively: it is cleat that the linear tails of log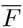(|*ξ*|) and logḠ(|*y*|) have the same slope −*α*.

Thus we can set, without loss of generality, *δt* = 1 and the discrete form of Eq. (2), the Euler-Maruyama equation [38], can be written simply as

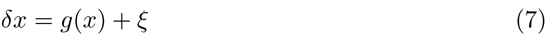

where *δx* is the finite time activity variation (cfr. Eq.(1) of the BOLD signal *x*(*t* + 1) – *x*(*t*)

The deterministic component can be easily recovered from the data by making use of condition (3); a computational procedure to calculate *g*(*x*) from measures of *x* at different times is presented in [39].

Samples of |*ξ*| can be obtained from the data once *g*(*x*) has been computed, via the formula

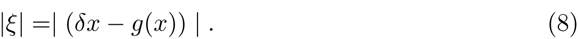

In principle, from these samples it is straightforward to derive the logarithm of the survival function, from which the parameter *α* can be fitted.

However, *α* is but one parameter of the distribution *ƒ*(*x;α,β,γ,δ*). It should be noted at this point that there is no closed-form formula for *ƒ*, which is often described by its characteristic function *E*[exp(itx)] = *∫*_ℝ_ exp(*itx*)*dF*(*x*), *F* being the cumulative distribution function (CDF). Explicitly,

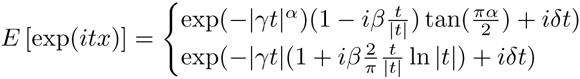

the first expression holding if *α* = 1, the second if *α* = 1. Special cases of stable distributions whose pdf can be written analytically, are given for *α* = 2, the normal distribution *ƒ*(*x*; 2, 0, *γ,δ*), for *α* = 1, the Cauchy distribution *ƒ*(*x*; 1,0,*γ,δ*), and for *α*= 0.5, the Lévy distribution *ƒ*(*x*;0.5,1,*γ, δ*); for all other cases, only the characteristic function is available in closed form, and numerical approximation techniques must be adopted for both sampling and parameter estimation [40-42].

Here, the method proposed in [42] has been used, which estimates the four parameters of an *α*-stable distribution via its characteristic function and was calculated by using Matlab implementation developed by M. Veillette (http://math.bu.edu/people/mveillet/html/alphastablepub.html).

The quality of the fit can then been assessed via the two-sample Kolmogorov-Smirnov (K-S) test, which is very sensitive in detecting statistical differences between two populations of data.

## Results

Calculation of *g*(*x*) for different networks have shown remarkably similar results, in that in any case the estimated *g*(*x*) can be fitted with a decreasing straight line of the form

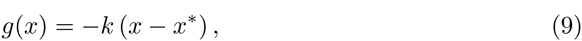

where *x*\* is defined by the condition *g*(*x*\*) = 0 and −*k* is the slope of the line; both *k* and *x*\* are specific for the network under consideration.

Graphs of *g*(*x*) for different networks are presented in Fig. 3 where for clarity’s sake all functions have been shifted so that *x*\* = 0.

**Figure 3.**
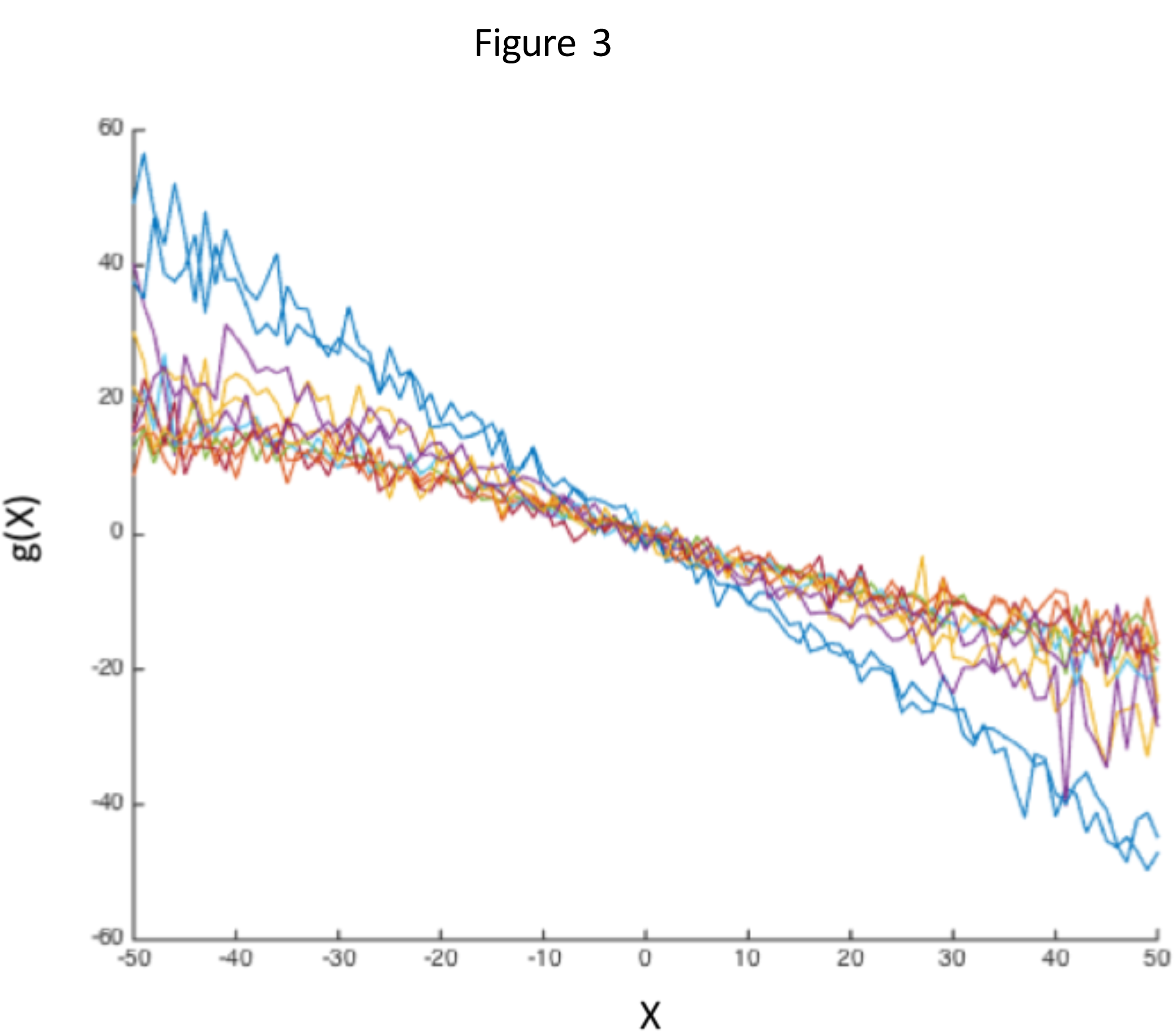
Graphs of the deterministic function *g*(*x*) for all networks. For clarity’s sake the points *x*\* for which *g*(*x*\*) = 0 have been shifted to 0. See text for explanations

Then Eq. (3), becomes

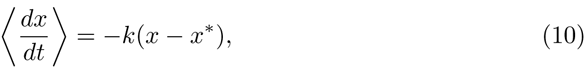

that can be easily solved to give 〈*x*〉 as function of time

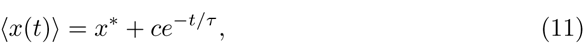

where *c* is a constant determined by the initial conditions and *τ* = 1/*k* is the characteristic time of the exponential decay.

Since (*x*) coincides with *x* when *ξ* = 0, the linearly decreasing form for the deterministic component has a simple meaning: it says that the activity *x*, in absence of the stochastic part, would tend exponentially to the equilibrium point *x*\*. Thus *g*(*x*) corresponds to the dissipation term in the Langevin equation for physical systems [15]

In contrast with the common form for *g* the distribution of stochastic term *ξ* appears to be peculiar of a given network, as revealed by the values of *α*, in general different acroos the networks.

This is apparent at a glance from the graphs of Fig. (4) where the empirical survival functions of amplitudes |*ξ*| are presented on a log-log scale together with the results of the fitting procedure.

**Figure 4.**
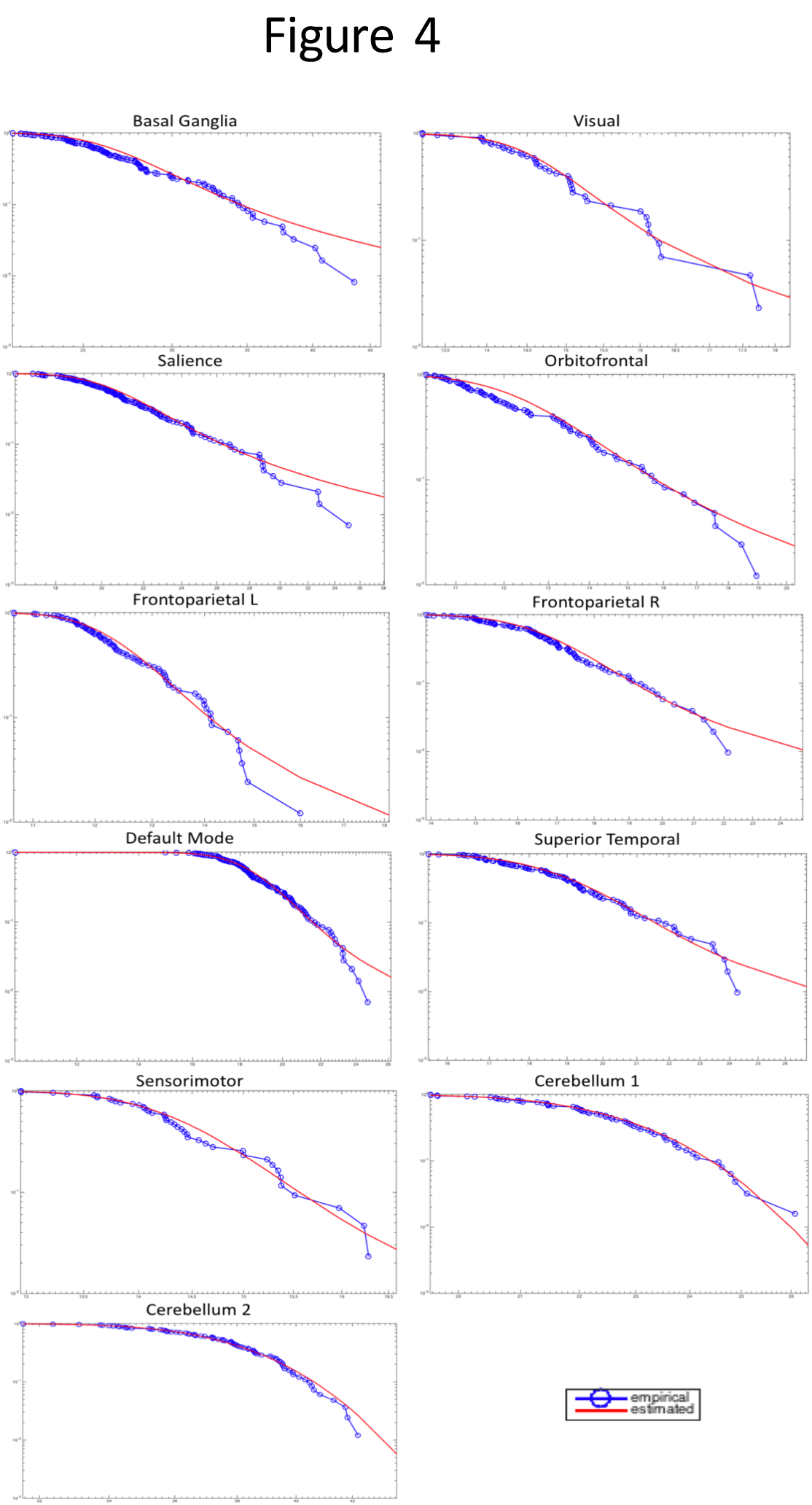
Comparison of logarithms of empirical and estimated survival function. Here a log-log plot has been used.

Statistical significance via the two-sample Kolmogorov-Smirnov (K-S) test (significance level *α* = 0.05) has shown that in all cases the null hypothesis *H*_0_ of no significant difference between the fit and the data cannot be rejected.

A summary of these findings can be found in Fig. (5) where values of *α* for different networks are represented in a bar graph, together with the corresponding brain areas.

**Figure 5.**
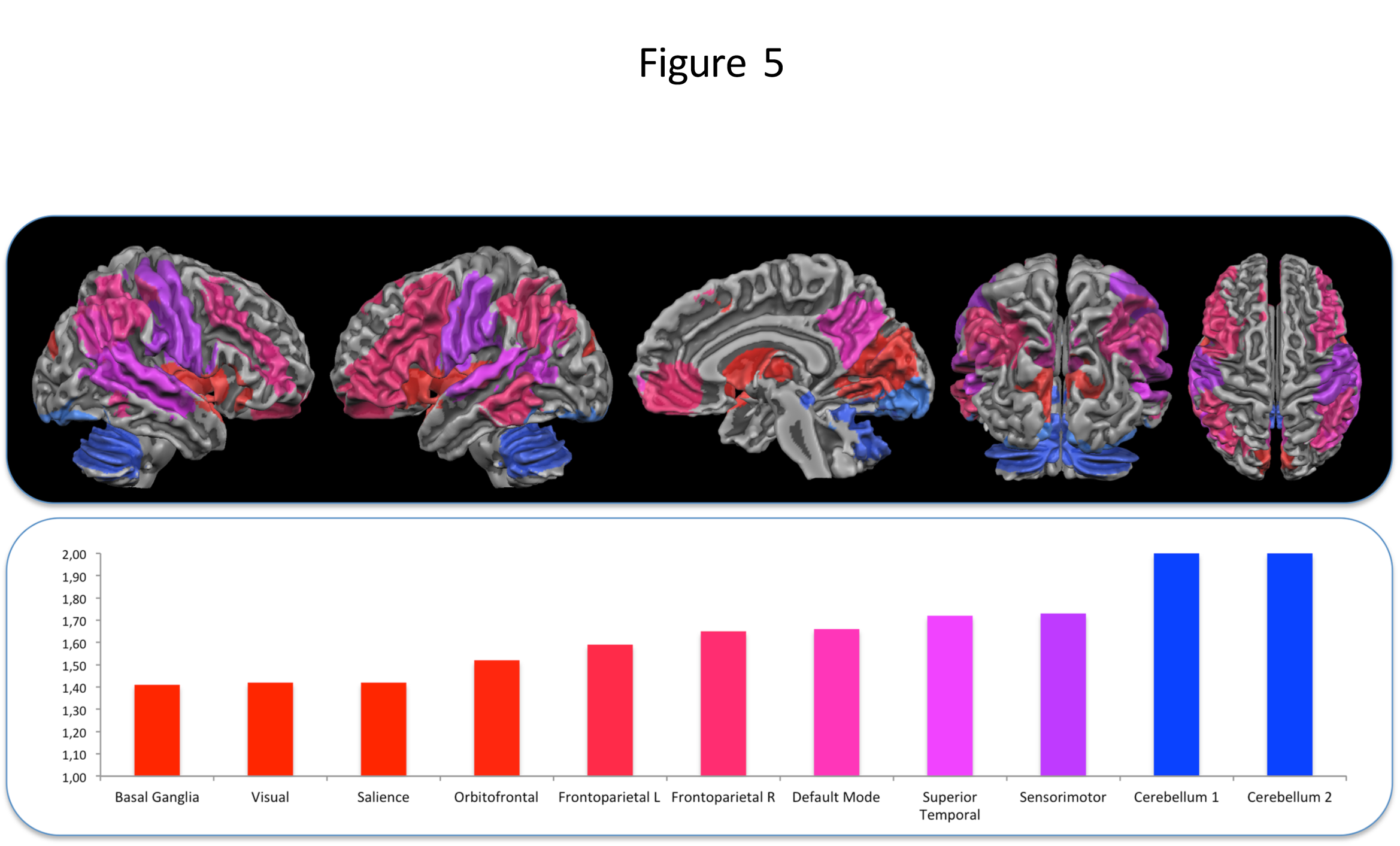
The graph represents the values of *α* for different networks from the lowest (red) to the highest. On the top of the graphs the corresponding brain areas are shown

The natural question arises of how activity patterns with different *α* values are related to distinct psychological states.

Here two maps have been considered, the first formed by the activity of the the three networks with the lowest *α* values and the second is derived from networks with the highest *α* values.

Discrimination of different functional meaning of the two maps has been carried out by making use of the Meta-Analytic Decoding of Network Function [43] that is a function of the Neurosynth framework [44]. Neurosynth databases contains several thousand psychological terms and topics and its built-in algorithms allows to draw inferences about the potential cognitive state associated with distributed patterns of activation.

For each map, we computed the voxel-wise Pearson correlation with each of 200 topic-based meta-analysis maps in the Neurosynth database (see http://neurosynth.org). The resulting coefficients were then used to generate a ranking of the psychological topics most consistently or specifically associated with each of the two maps.

The results of this procedure are shown as a “radar map” of FIg. 6, where the most relevant categories associated with the activity patterns are shown at positions determined by the corresponding correlation coefficients.

**Figure 6.**
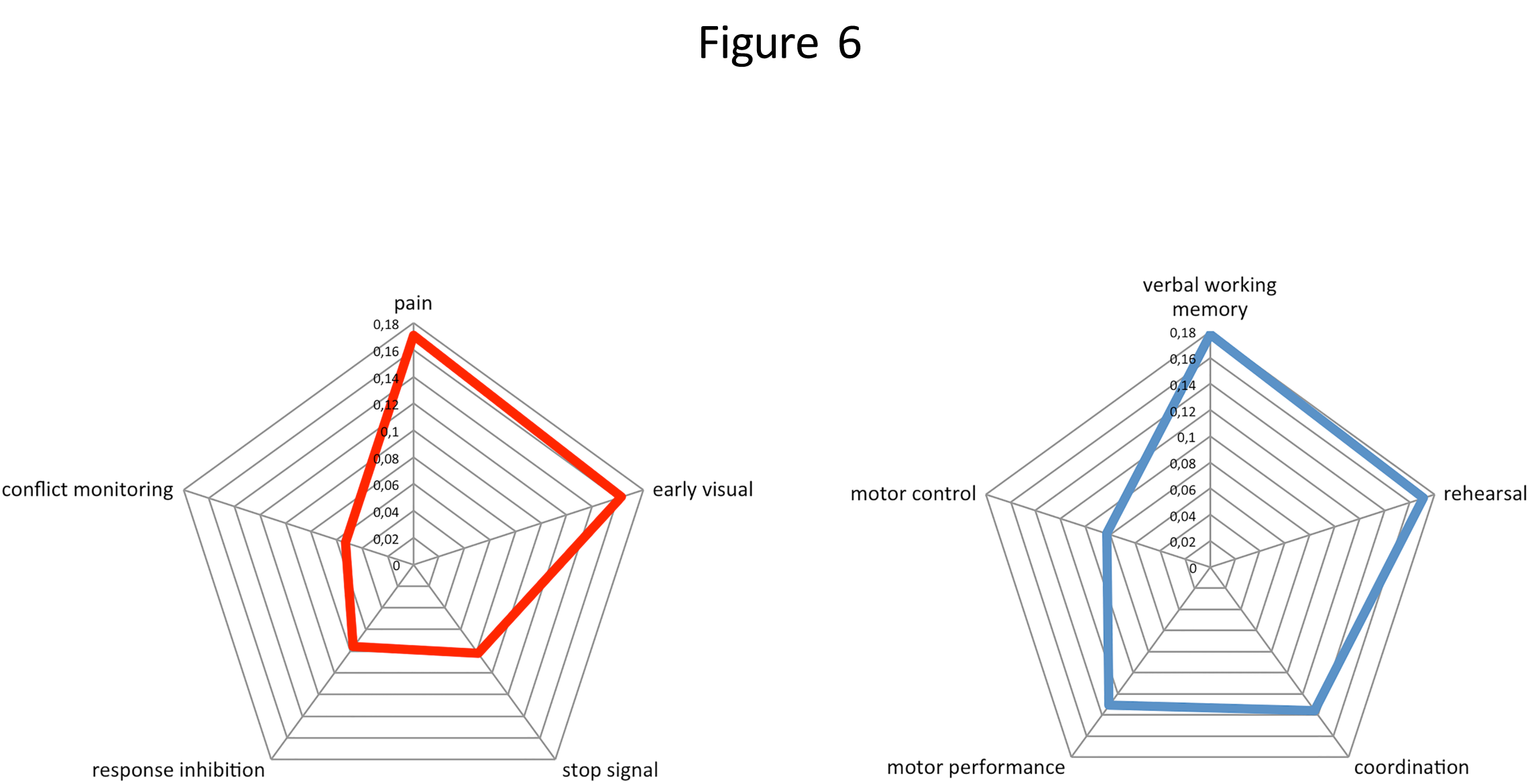
“Radar map” representing the most relevant cognitive terms for two maps corresponding to low (red line) and high *α* values (blue line)

The relevant states for the low *α* map appear to represent categories related to sensory processes whereas the high *α* map have more to do with motor control and routines‥

## Discussion

In this work we have investigated the variation or activity of the BOLD signal relying upon the stable Lévy process. We have found that activity in different networks of the brain can be explained by resorting to a generalized Langevin stochastic differential equation, with a simple additive interaction between the deterministic drift term and the stochastic term, namely *α*-stable noise.

All networks share the same deterministic part, namely an exponentially decreasing relaxation term to some steady state of activity, the only difference being the values of these equilibrium points.

This relaxation term is common to many models of neural networks [45], [33], and, as mentioned before, has a simple meaning; after all it is natural for the activity to reach some equilibrium level. This also shows that most relevant information of the signal is contained in the random term.

For all networks the stochastic variable *ξ* follows an *α*-stable distribution, with, in general, different values of *α*.

In particular, the analysis has demonstrated the existence of a range of values of *α*, and therefore of dynamics: for instance, basal ganglia, visual and salience networks seems present a definite “Lévy motion”-like behavior of activity, whereas areas from the cerebellum exhibit “Brownian motion” (i.e., *α* = 2, originating Gaussian dynamics).

In the “Brownian motion” case fluctuations of x are small and spread relatively evenly among a range of possible values, whilst “Lévy motion” presents a dynamics in which variations are usually very small, close to 0, but where large fluctuations are more likely than in the Gaussian case.

Such difference in behaviors is relevant from a biological signal processing point of view: Gaussian dynamics is consistent with some form of continuous coding, whereas, by contrast a Lévy dynamics can be considered to implement a sparse coding in time via a subordination mechanism [46].

Indeed, in sparse coding theory activity distributions are assumed to be highly peaked around zero and with heavy tails, which is the case also for displacements in Lévy flights and, moreover, in many computational models activity is assumed to follows, a priori, a Cauchy distribution, which is also the most used to generate Lévy flights for a variety of applications, indeed it can be considered a paradigmatic Lévy distribution.

Finally, theoretical studies and empirical evidence suggest that visual cortex uses a sparse code to efficiently represent natural scenes [47] and this is consistent with the evidence, presented here, that activity of the “visual” area follows a Levy distribution. What are the reasons by which these specific networks have these basically different behaviours?

It should be noted that Lévy flights can be found in networks more involved with interacting with the external world, and a reason may be that large fluctuations are more efficient in exploring the activity space. Optimality of Lévy flights has been proved for a variety of searches from those related to animal foraging [48,49] and human mobility [50] to visual information foraging based on eye movements [25-28]. Interestingly enough, it has been conjectured that what was once foraging for tangible resources in a physical space has adapted to foraging in cognitive space for information related to those resources (e.g., goal-directed deployment of visual attention), with a fundamental role played by the dopaminergic control [30]. On the other hand a recent study [14] has pointed out how BOLD variability is related to dopaminergic neurotransmission. Our results are in line with these studies, indeed the main dopaminergic area has the smallest *α* value, among the areas considered here.

Another point worth highlighting is that areas with *α* = 2 belong to the cerebellum; thus activity in the cerebellum appears to be driven by a Gaussian dynamics. Doya [51] has hypothesized that the cerebellum consists of a number of independent modules, all with the same internal structure and performing the same computation, depending on the input-output connections of the module. The cerebellum has been implicated in the regulation of many differing functional traits such as affection, emotion and behavior. This type of regulation can be understood as predictive action selection based on internal models of the environment and it is well known that a Gaussian process can be used as a prior probability distribution over functions in Bayesian inference.

Further support for these suggestions derives from the results of Meta-Analytic Decoding of Network Function showing that maps corresponding to the networks with an *α* ≈ 2 are involved in cognitive function like coordination and motor performance, on the other hand the map with the three networks with the lower value of *α* are involved in cognitive function like early visual, response inhibition and pain that are all cognitive functions requiring with an high level of attention. Then it is possible to hypothesize that a more rapid variation of the signal is needed to cover more states in a given time interval.

Finally, results of this work can be useful to investigate dynamics of the brain activity in pathological conditions or during aging: in particular one can look to variations of the *α* to discriminate between normal and pathological conditions.

## References

1. Deco G, Jirsa VK, McIntosh AR. Emerging concepts for the dynamical organization of resting-state activity in the brain. Nature Reviews Neuroscience. 2011;12(1):43–56.

2. Papo D. Functional significance of complex fluctuations in brain activity: from resting state to cognitive neuroscience. Frontiers in Systems Neuroscience. 2014;8(112).

3. Friston KJ, Holmes AP, Worsley KJ, Poline JP, Frith CD, Frackowiak RS. Statistical parametric maps in functional imaging: a general linear approach. Human brain mapping. 1994;2(4):189–210.

4. Haxby JV, Gobbini MI, Furey ML, Ishai A, Schouten JL, Pietrini P. Distributed and overlapping representations of faces and objects in ventral temporal cortex. Science. 2001;293(5539):2425–2430.

5. Mourao-Miranda J, Bokde AL, Born C, Hampel H, Stetter M. Classifying brain states and determining the discriminating activation patterns: Support Vector Machine on functional MRI data. NeuroImage. 2005;28(4):980–995.

6. Pereira F, Mitchell T, Botvinick M. Machine learning classifiers and fMRI: a tutorial overview. Neuroimage. 2009;45(1):S199–S209.

7. Fox MD, Raichle ME. Spontaneous fluctuations in brain activity observed with functional magnetic resonance imaging. Nature Reviews Neuroscience. 2007;8(9):700–711.

8. Roebroeck A, Formisano E, Goebel R. Mapping directed influence over the brain using Granger causality and fMRI. Neuroimage. 2005;25(1):230–242.

9. Friston KJ. Modalities, modes, and models in functional neuroimaging. Science. 2009;326(5951):399–403.

10. Bianciardi M, Fukunaga M, van Gelderen P, Horovitz SG, de Zwart JA, Duyn JH. Modulation of spontaneous fMRI activity in human visual cortex by behavioral state. Neuroimage. 2009;45(1):160–168.

11. Grady CL, Garrett DD. Understanding variability in the BOLD signal and why it matters for aging. Brain imaging and behavior. 2014;8(2):274–283.

12. He BJ. Scale-free properties of the functional magnetic resonance imaging signal during rest and task. The Journal of neuroscience. 2011;31(39):13786–13795.

13. Garrett DD, Samanez-Larkin GR, MacDonald SW, Lindenberger U, McIntosh AR, Grady CL. Moment-to-moment brain signal variability: a next frontier in human brain mapping? Neuroscience & Biobehavioral Reviews. 2013;37(4):610–624.

14. Guitart-Masip M, Salami A, Garrett D, Rieckmann A, Lindenberger U, Bäckman L. BOLD variability is related to dopaminergic neurotransmission and cognitive aging. Cerebral Cortex. 2015;p. bhv029.

15. Risken H. Fokker-Planck Equation. Springer; 1984.

16. Van Kampen NG. Stochastic processes in physics and chemistry. Amsterdam, NL: North Holland; 2001.

17. Dubkov AA, Spagnolo B, Uchaikin VV. Lévy flight superdiffusion: an introduction. International Journal of Bifurcation and Chaos. 2008;18(09):2649–2672.

18. Bouchaud JP, Georges A. Anomalous diffusion in disordered media: statistical mechanisms, models and physical applications. Physics reports. 1990;195(4):127–293.

19. Metzler R, Klafter J. The random walk’s guide to anomalous diffusion: a fractional dynamics approach. Physics reports. 2000;339(1):1–77.

20. Posadas A, Morales J, Vidal F, Sotolongo-Costa O, Antoranz J. Continuous time random walks and south Spain seismic series. Journal of seismology. 2002;6(1):61–67.

21. Janssen H, Oerding K, Van Wijland F, Hilhorst H. Lévy-flight spreading of epidemic processes leading to percolating clusters. The European Physical Journal B-Condensed Matter and Complex Systems. 1999;7(1):137–145.

22. Scafetta N, Latora V, Grigolini P. Lévy scaling: the diffusion entropy analysis applied to DNA sequences. Physical Review E. 2002;66(3):031906.

23. Viswanathan GM, Afanasyev V, Buldyrev S, Murphy E, Prince P, Stanley HE, et al. Lévy flight search patterns of wandering albatrosses. Nature. 1996;381(6581):413–415.

24. Rhodes T, Turvey MT. Human memory retrieval as Lévy foraging. Physica A: Statistical Mechanics and its Applications. 2007;385(1):255–260.

25. Brockmann D, Geisel T. The ecology of gaze shifts. Neurocomputing. 2000;32(1):643–650.

26. Boccignone G, Ferraro M. Modelling gaze shift as a constrained random walk. Physica A. 2004;331:207–218.

27. Boccignone G, Ferraro M. Ecological Sampling of Gaze Shifts. IEEE Trans on Cybernetics. 2014 Feb;44(2):266–279.

28. Marlow CA, Viskontas IV, Matlin A, Boydston C, Boxer A, Taylor RP. Temporal Structure of Human Gaze Dynamics Is Invariant During Free Viewing. PloS one. 2015;10(9):e0139379.

29. Tsallis C, Levy SV, Souza AM, Maynard R. Statistical-mechanical foundation of the ubiquity of Lévy distributions in nature. Physical Review Letters. 1995;75(20):3589.

30. Hills TT. Animal Foraging and the Evolution of Goal-Directed Cognition. Cognitive Science. 2006;30(1):3–41.

31. Gerstein GL, Mandelbrot B. Random walk models for the spike activity of a single neuron. Biophysical journal. 1964;4(1 Pt 1):41.

32. Capocelli R, Ricciardi L. Diffusion approximation and first passage time problem for a model neuron. Kybernetik. 1971;8(6):214–223.

33. Linaro D, Storace M, Giugliano M. Accurate and fast simulation of channel noise in conductance-based model neurons by diffusion approximation. PLoS Comput Biol. 2011;7(3):e1001102.

34. West BJ. Fractal physiology and the fractional calculus: a perspective. Frontiers in physiology. 2010;1.

35. Jespersen S, Metzler R, Fogedby HC. Lévy flights in external force fields: Langevin and fractional Fokker-Planck equations and their solutions. Physical Review E. 1999;59(3):2736.

36. Van Kampen NG. Stochastic processes in physics and chemistry. vol. 1. Elsevier; 1992.

37. Gnedenko BV, Kolmogorov AN, Chung KL, Doob JL. Limit distributions for sums of independent random variables. 1954;.

38. Higham DJ. An algorithmic introduction to numerical simulation of stochastic differential equations. SIAM review. 2001;43(3):525–546.

39. Siegert S, Friedrich R. Modeling of nonlinear Lévy processes by data analysis. Physical Review E. 2001;64(4):041107.

40. Chambers J, Mallows C, Stuck B. A method for simulating stable random variables. J Am Stat Ass. 1976;71(354):340–344.

41. Nolan JP. Numerical calculation of stable densities and distribution functions. Communications in Statistics-Stochastic Models. 1997;13(4):759–774.

42. Koutrouvelis IA. An iterative procedure for the estimation of the parameters of stable laws: An iterative procedure for the estimation. Communications in Statistics-Simulation and Computation. 1981;10(1):17–28.

43. Chang LJ, Yarkoni T, Khaw MW, Sanfey AG. Decoding the role of the insula in human cognition: functional parcellation and large-scale reverse inference. Cerebral Cortex. 2013;23(3):739–749.

44. Yarkoni T, Poldrack RA, Nichols TE, Van Essen DC, Wager TD. Large-scale automated synthesis of human functional neuroimaging data. Nature methods. 2011;8(8):665–670.

45. Wilson HR. Spikes, decisions, and actions: the dynamical foundations of neuroscience. Oxford University Press; 1999.

46. Brockmann D, Sokolov I. Lévy flights in external force fields: from models to equations. Chemical Physics. 2002;284(1):409–421.

47. Olshausen BA, Field DJ. Natural image statistics and efficient coding. Network: computation in neural systems. 1996;7(2):333–339.

48. Méndez V, Campos D, Bartumeus F. Stochastic foundations in movement ecology: anomalous diffusion, front propagation and random searches. Springer Science & Business Media; 2013.

49. Viswanathan GM, Da Luz MG, Raposo EP, Stanley HE. The physics of foraging: an introduction to random searches and biological encounters. Cambridge University Press; 2011.

50. Brockmann D, Hufnagel L, Geisel T. The scaling laws of human travel. Nature. 2006;439(7075):462–465.

51. Doya K. Complementary roles of basal ganglia and cerebellum in learning and motor control. Current opinion in neurobiology. 2000;10(6):732–739.

